# *In vitro* evidence for bisphenol A as a human liver carcinogen: Environmentally relevant doses inhibit cancer-protective ESR1 signaling in a human liver cell line

**DOI:** 10.64898/2025.12.18.695265

**Authors:** Emma Weeks, Scott Kennedy, Robert Searles, Samantha Carrothers, Brett Davis, Lucia Carbone, Mitchell Turker, R. Stephen Lloyd, Caren Weinhouse

## Abstract

Several environmentally ubiquitous endocrine disrupting chemicals (EDCs) are suspected carcinogens, but their mechanism(s) of action are unknown. In this study, we test the potential for a model EDC, bisphenol A (BPA), to both initiate (via oxidative mutagenesis) and promote (via endocrine disruption) liver carcinogenesis. This study is motivated by our prior finding that developmental BPA exposure caused hepatocellular carcinoma (HCC) in mice. Here, we provide *in vitro* evidence supporting a mechanism for BPA as a non-genotoxic carcinogen. Using a highly sensitive, error-corrected sequencing method, we demonstrate that human population-relevant doses of BPA cause mutations that are consistent with oxidative DNA damage; however, overall mutation frequency does not differ substantially from controls. In contrast, we show that BPA inhibits cancer-protective, estrogen-induced transcription of estrogen receptor 1 (ESR1) target genes in the presence of pre-pubertal, but not post-pubertal, levels of estradiol. These results constitute strong initial evidence supporting BPA as a liver cancer promoting agent. This mechanism may be generalizable to a wide range of environmental EDCs that are weak agonists for ESR1. This finding is critically important to prevention of HCC, which is prevalent, lethal, and poorly responsive to therapy.

## II. Introduction

Endocrine disrupting chemicals (EDCs), a class of synthetic compounds that disrupt hormone signaling, are controversial carcinogens^2-8^. EDC exposure to the U.S. population is ubiquitous and chronic^9,10^, so any adverse outcomes of these exposures are likely to have large impacts on population health. EDCs’ carcinogenicity is unclear largely because of our limited understanding of their carcinogenic mechanism(s) of action^3,11^. Most carcinogens are classified as genotoxic or non-genotoxic^12,13^. Genotoxic compounds cause cancer by damaging DNA. If unrepaired, this damage leads to DNA mutations; high mutation loads increase the probability of driver mutations occurring in oncogenes or tumor suppressor genes that initiate cancer^12,13^. In contrast, non-genotoxic carcinogens cause cancer via promotion, or stimulation of proliferation of initiated cells carrying driver mutations^14^. *In vitro* studies of many EDCs show their ability to damage DNA, often as a secondary effect of increased cellular reactive oxygen species (ROS)^2,15^. However, these studies employ doses several orders of magnitude higher than common human exposures^15^, which leaves open the question of whether environmentally relevant doses cause similar levels of DNA damage. In addition, it is unclear whether EDCs induce sufficient DNA damage to overwhelm repair mechanisms and lead to accumulating DNA mutations that can drive cancer. Evidence for non-genotoxic effects of EDCs is stronger. Specifically, both *in vitro* and *in vivo* evidence supports EDC-induced cell proliferation, often stimulated through nuclear hormone receptor signaling^11,16^. However, few studies have shown convincingly that this proliferative effect is sufficient to promote cancer development.

In this study, we test the carcinogenic potential of a model EDC, bisphenol A (BPA), in a human liver cancer cell line, HepG2. This focus is motivated by our prior finding that developmental exposure to environmentally relevant doses of BPA increases the incidence of hepatocellular carcinoma (HCC) in C57BL/6J mice^4^. In our design, we address three main limitations of prior studies of BPA’s carcinogenicity. First, we use low doses that mimic human exposure levels. Second, we focus on the liver, which is a target organ of endogenous hormones that is commonly overlooked in favor of better-known targets, primarily reproductive tract organs (e.g., prostate, testes, breast, ovaries)^16,17^. Third, we test proposed mechanisms for both cancer initiation and promotion by BPA.

The design of this study is based on four characteristics of BPA-induced HCC that we observed in our prior work^4^. These four characteristics are as follows. First, we observed a dose-responsive relationship, in which HCC rates were higher in mice exposed to higher doses of BPA^4^, which is characteristic of genotoxic initiation^12,18^. Second, we observed that HCC developed at similar rates in both male and female mice^4^, which is notable, since human and rodent males are 2-4x more likely to develop HCC, as compared to females^19-21^. This male bias holds true for all known HCC initiating and promoting agents in rodents and humans, including chemical carcinogens aflatoxin B_1_ (AFB_1_)^22^, diethyl nitrosamine (DEN)^19,23^ and phenobarbital^19,23^, as well as non-chemical cancer risk factors, including hepatitis B^20,24,25^ or C^20,26^ infection and alcohol abuse. This sex bias is due to protective estrogen receptor 1 (ESR1) signaling in females^19^. Therefore, our observation suggests that BPA promotes HCC in females through disruption of this protective mechanism. Third, we observed cancer in mice exposed during development to low doses of BPA^4^, despite earlier reports of no cancer in either Fischer 344 rats or B6C3F1 hybrid mice following high doses of BPA in adulthood^27^. This result suggests a developmental susceptibility to BPA carcinogenesis. (We note here that the recent CLARITY-BPA study reported no liver cancer in Sprague-Dawley rats (NCTR strain) exposed during development to low dose BPA^17^. However, this rat strain’s lack of responsiveness to the positive control ethinyl estradiol^28^ demonstrates that it lacks utility in assessing risk of BPA exposure^16,17,29^.) Fourth, we observed liver cancer in mice exposed only to BPA, with no known co-exposures, suggesting that BPA acts as a complete carcinogen in liver.

A complete carcinogen is capable of both initiating and promoting cancer. If BPA can initiate liver cancer, it most likely does so by causing oxidative DNA damage at high enough levels to cause increased mutation loads. The evidence linking BPA and oxidative stress is strong. High-dose BPA increases cellular ROS *in vitro* and *in vivo*, and epidemiologic studies have linked BPA exposure to markers of oxidative stress in humans^15^. A smaller number of studies have shown that BPA-induced ROS leads to oxidatively induced damage to DNA^30,31^. However, only one study to date has shown that the DNA damage caused by high dose BPA leads to increased mutation load^15,32^; therefore, it remains an open question whether low dose BPA is mutagenic. However, if BPA is mutagenic, the liver is likely most susceptible to its effects, because the liver is exposed to the highest dose of bioactive BPA due to first-pass metabolism^11,16,17^.

If BPA can promote liver cancer, it most likely does so via disruption of protective ESR1 signaling. Broadly, ESR1 regulates two gene expression programs in liver: a mitogenic program that promotes cellular proliferation, which increases cancer risk, and a protective program that limits cellular proliferation^19^. The first program is controlled by enhancers bound by ESR1 alone; the second is regulated by enhancers co-bound by forkhead box A (FoxA) factors, FoxA1 and FoxA2^19^. In healthy females, the protective program dominates the proliferative one^19^. However, when ESR1 cannot bind FoxA enhancers, females develop more HCC, due to the loss of the protective program and the resulting dominance of the proliferative program^19^. Therefore, BPA likely disrupts the protective program. This is biologically feasible; when liganded by BPA, ESR1 is likelier to bind preferentially to FoxA enhancers, and, once bound, to inhibit transcription of protective genes. BPA is a weak ESR1 agonist and forms a complex with the receptor^33,34^ that requires cooperative binding by other transcription factors, like FoxA factors, in order to bind DNA^34^. In addition, the BPA-ESR1 complex is less capable of recruiting transcriptional co-factors^33-35^, as compared to the E_2_-ESR1 complex. Therefore, BPA-ESR1 binding does not trigger transcriptional change at target genes^34,35^. Taken together, our working model is that BPA inhibits the transcriptional activity of ESR1 at FoxA enhancers, thereby inhibiting the HCC protective program in females. It is also feasible that these effects are limited to the pre-pubertal window. BPA has substantially lower affinity for ESR1 than E_2_^33,35^ and is outcompeted in the presence of high E_2_ (e.g., in post-pubertal females)^34^. Therefore, developmental, but not adult, BPA exposure is most likely to increase liver cancer risk.

Here, we test whether low dose BPA exposure supports either or both of our proposed mechanisms. Specifically, we test whether BPA induces mutations that reflect oxidative DNA damage and inhibits ESR1 signaling in the presence of developmental, but not adult, levels of E_2_ (**Fig. 1**). We show strong evidence for ESR1 inhibition by low dose BPA at developmental levels of E_2_, which supports a role for developmental BPA in liver cancer promotion. We show that low dose BPA causes mutations that are consistent with oxidative DNA damage; however, overall mutation frequency does not differ substantially from controls. These results provide strong supporting evidence for BPA as a non-genotoxic carcinogen, when exposure occurs prior to pubertal onset.

**Figure 1.**
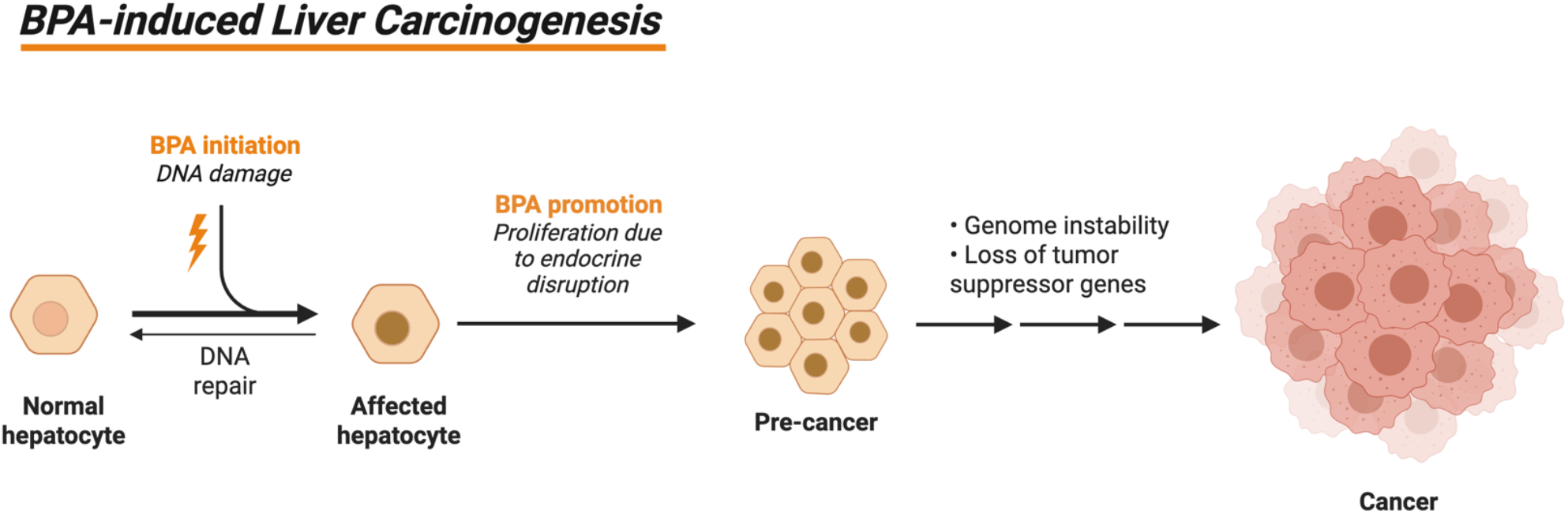
Proposed model of BPA-induced liver carcinogenesis. To function as a complete carcinogen, BPA must both initiate and promote cancer development. BPA initiates cancer through DNA damage and mutagenesis. BPA then promotes cancer through cellular proliferation due to endocrine disruption. If initiation overwhelms DNA repair and promotion evades apoptosis, then pre-cancerous cells progress to cancer.

## III. Results

### Characterization of BPA-induced mutations via Duplex-seq

First, we tested our hypothesis that BPA can initiate cancer via unrepaired oxidatively induced DNA damage that leads to driver mutations (**Fig. 2A**). To detect mutations induced by low dose BPA exposure, we leveraged a powerful next-generation sequencing method called error-corrected Duplex sequencing^13,36^. The background error rate of this method is ∼2×10^-8^, which is 100-1,000X lower than the background frequency of somatic mutations and >10,000X below conventional next-generation sequencing methods^13,36^. Real world doses of environmental chemicals often cause low level mutations that are below the limit of detection of less sensitive methods; however, these mutations can be sufficient to initiate cancer, making their detection a critical missing link between chemical exposure and subsequent cancer^13^.

**Figure 2.**
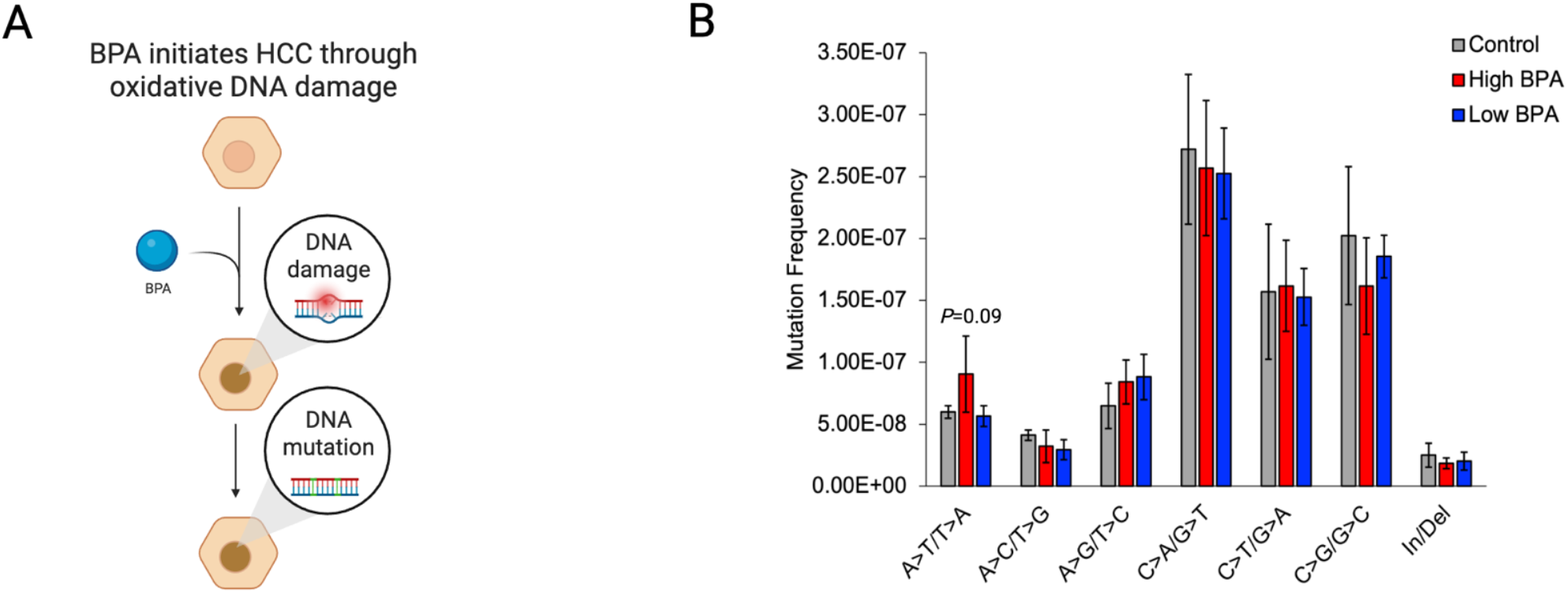
BPA exposure increases oxidatively induced DNA damage in HepG2 cells. **A**. Proposed model for initiation of HCC by BPA. **B**. Total mutation frequency detected by Duplex sequencing mutations in HepG2 cells exposed to DMSO (vehicle control) or BPA (0.4 μM or 0.04 μM) (*N*=4 per group) for 24h and sequenced with Duplex sequencing 48h post-exposure.

To perform these experiments, we exposed HepG2 cells for 24 hours to two doses of BPA (0.4 μM or 0.04 μM). These doses are the *in vitro* equivalent of liver exposure following *in vivo* exposure to 5 μg or 0.5 μg /kg BW/day (calculated using the NTP Integrated Chemical Environment PBPK tool), reflecting the maximum and median human population exposure levels^37^. We cultured cells for 48 hours post-exposure, to allow for mutation fixation^38^. Next, using DNA extracted from these cells, we prepared Duplex sequencing libraries that span 48 Kb of DNA targeting 20 genomic regions target 48 Kb of DNA across 20 genomic regions^13^. We sequenced these libraries to ∼15,000X and compared mutation frequencies and spectra between groups using a custom bioinformatics pipeline (see Methods). We observed a marginal increase in overall mutation load (ANOVA p=0.09), largely driven by an increase in A>T/T>A transversions (p=0.09 on Student’s t-test) (**Fig. 2B**) We noted an apparent increase in A>G/T>C transitions (**Fig. 2B**), despite not reaching a statistical threshold. Next, we screened for mutation signatures curated in the Catalogue of Somatic Mutations in Cancer (COSMIC) database and detected a single-base substitution (SBS) signature, SBS8, of unknown etiology (**Supp. Fig. S1**). We did not detect any double-base substitution (DBS) signatures (**Supp. Fig. S2**).

These results are consistent with some of the DNA base lesions previously observed in response to high dose BPA exposures. For example, high dose BPA exposure increased rates of the oxidized base lesion 5,6-dihydroxy-5,6-dihydropyrimidine, or thymine glycol^31^; unrepaired thymine glycol lesions cause A>G/T>C mutations^39^. In the presence of a second oxidizing agent, high dose BPA also increased rates of oxidized base lesions 4,6-diamino-5-formamido-adenosine^31^, or FapyAde; unrepaired FapyAde lesions cause A>T/T>A mutations^40^. In addition, five COSMIC SBS mutation signatures have been reported previously for BPA, including SBS8^32^, which we observed in this study.

However, our results do not support all prior data supporting BPA as an oxidative mutagen. We did not detect classic signatures of ROS-induced mutations, including G>T/C>A transversions, which reflect oxidatively induced 2,6-diamino-4-hydroxy-5-formamido-guanosine (FapyGua)^41^ and 8-oxo-2’-deoxyguanosine (8-oxo-dG) base lesions^42^, as well as bulky adducts formed by BPA quinone metabolites^43-47^. Neither did we detect the COSMIC signature SBS18 (“possibly damage by reactive oxygen species”)^48,49^. In addition, we did not observe small indels with microhomology reflective of DNA double-strand breaks (DSBs) (COSMIC signature ID6)^48^, although ROS-induced DSBs have been reported in BPA-exposed germ cells in multiple prior studies^50-52^.

### Evaluation of ESR1-FOXA1/2 target gene expression following BPA exposure

To test our hypothesis that ESR1 target gene expression would be inhibited by BPA in the presence of low but not high E_2_ (**Fig. 3A**), we exposed HepG2 cells to six different exposures for 24 hours, following five-day pre-incubation in charcoal-stripped fetal bovine serum and either vehicle (DMSO), low E_2_, or high E_2_, to mimic the group-relevant background hormonal environment: 1) 0.4 μM BPA; 2) 5 pM E_2_ (equivalent to circulating levels in pre-pubertal females^34^); 3) 10 nM E_2_ (equivalent to circulating levels in post-pubertal females^34^); 4) 0.4 μM BPA + 5 pM E_2_ (modeling an exposed pre-pubertal female); 5) 0.4 μM BPA + 10 nM E_2_ (modeling an exposed post-pubertal female); or 6) DMSO vehicle control.

**Figure 3.**
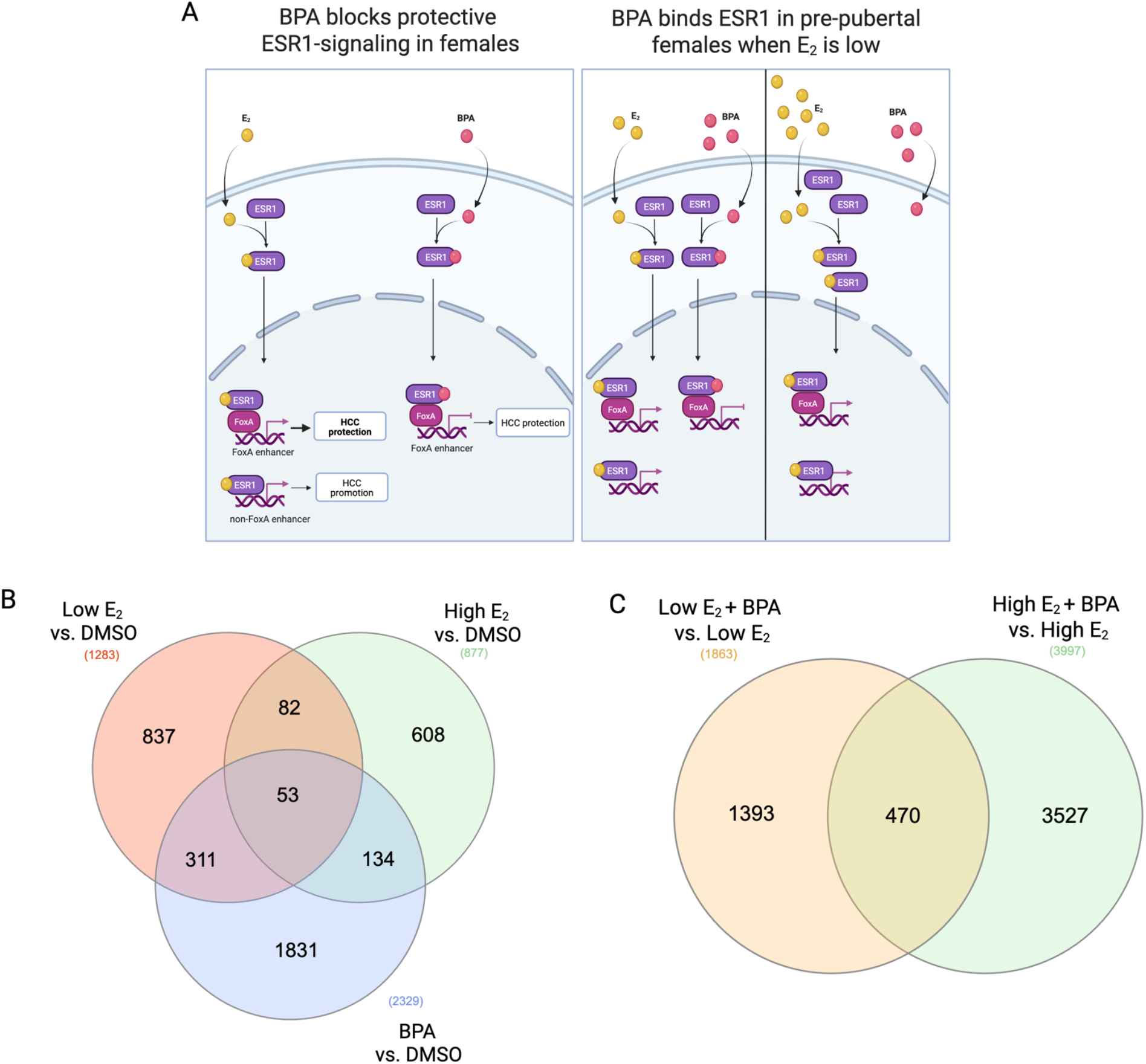
BPA exposure disrupts gene expression in HepG2 cells with physiologically relevant circulating estrogen levels. **A**. Proposed model of BPA-induced HCC promotion. **B-C**. Differentially expressed genes (DEGs) in HepG2 cells exposed to: (B) low E_2_ (5 pM; equivalent to pre-pubertal circulating levels), high E_2_ (10 nM; equivalent to post-pubertal circulating levels), or BPA (0.4 μM; equivalent to the liver dose of common human population exposure), compared to DMSO vehicle control; or (C) low E_2_ + BPA (compared to low E_2_ alone; models exposed vs. unexposed pre-pubertal females) or high E_2_ + BPA (compared to high E_2_ alone; models exposed vs. unexposed post-pubertal females).

First, we identified differentially expressed genes (DEGs) in each pair-wise comparison. Using a stringent false discovery rate (FDR) cutoff of 0.05, to account for false positives attributable to multiple statistical tests, we identified between 12 and 886 DEGs in pair-wise comparisons. We attribute this limited result to within-group variation among replicates in some experimental groups (**Supp. Fig. S3**), possibly due to our focus on environmentally relevant low doses of BPA and physiologically relevant low doses of E_2_. To maximize visibility of informative patterns in our dataset, we performed subsequent steps using DEGs with p<0.05, without multiple testing correction. All reported patterns held true when limited to genes with FDR<0.05. We identified 53 genes (fewer than 3%) that were differentially expressed in all three, single exposure comparisons (low E_2_ vs. DMSO, high E_2_ vs. DMSO, BPA vs. DMSO); the majority of DEGs were unique to each comparison group (**Fig. 3B**). In contrast, we identified 470 DEGs (12-25%) in common between the two groups designed to mimic human exposures: low E_2_ + BPA vs. low E_2_ (which models BPA-exposed vs. -unexposed pre-pubertal females) and high E_2_ + BPA vs. high E_2_ (which models BPA-exposed vs. -unexposed post-pubertal females) (**Fig. 3C**). Gene Ontology (GO) enrichment analyses for these last two exposure groups showed general enrichment for critical cellular processes, including protein localization to cellular membranes and subcellular compartments and synthesis of cellular components (**Supp. Fig. 4**). Notably, we observed several GO terms associated with cellular stress and cell proliferation in the high E_2_ + BPA vs. high E_2_ group (e.g., cellular response to stress, cellular catabolic process, mitotic cell cycle) that were not present in the low E_2_ + BPA vs. low E_2_ group (**Supp. Fig. 4**).

Next, we tested whether DEGs were enriched for genes co-regulated by ESR1 and FOXA factors. We filtered DEGs from the two experimental groups designed to model human exposure for genes within 10 Kb of a published ESR1 ChIP-seq peak from healthy human hepatocytes^1^ (to avoid aberrant binding events in a cancer cell line) and within 50 Kb of a FOXA1/2 sequence motif. In each comparison, approximately half of the DEGs detected met our filtering criteria (N=938/1863 genes in low E_2_ + BPA vs. low E_2_; N=2039/3997 genes in high E_2_ + BPA vs. high E_2_) (**Fig. 4A**); nearly all genes near ESR1 peaks were also near FOXA motifs and most that did not meet both criteria were located on the mitochondrial genome. The ESR1 ChIP-seq datasets^1^ that we used reported on ESR1 binding in human hepatocytes with or without E_2_ treatment; data were reported as “E2-treated”, which includes all ESR1 ChIP-seq peaks in E_2_-treated hepatocytes, or “E2-gained peaks”, which includes only ESR1 ChIP-seq peaks that were newly gained in the E_2_-treated condition but were not present without E_2_ treatment. We filtered our DEGs using both ChIP-seq datasets; in each case, all genes associated with “E2-gained peaks” were fully embedded in the set of genes associated with “E2-treated peaks” (**Fig. 4B-C**) and all observed patterns held true in both gene sets. Therefore, we focus on results from the larger set of genes associated with “E2-treated peaks.”

**Figure 4.**
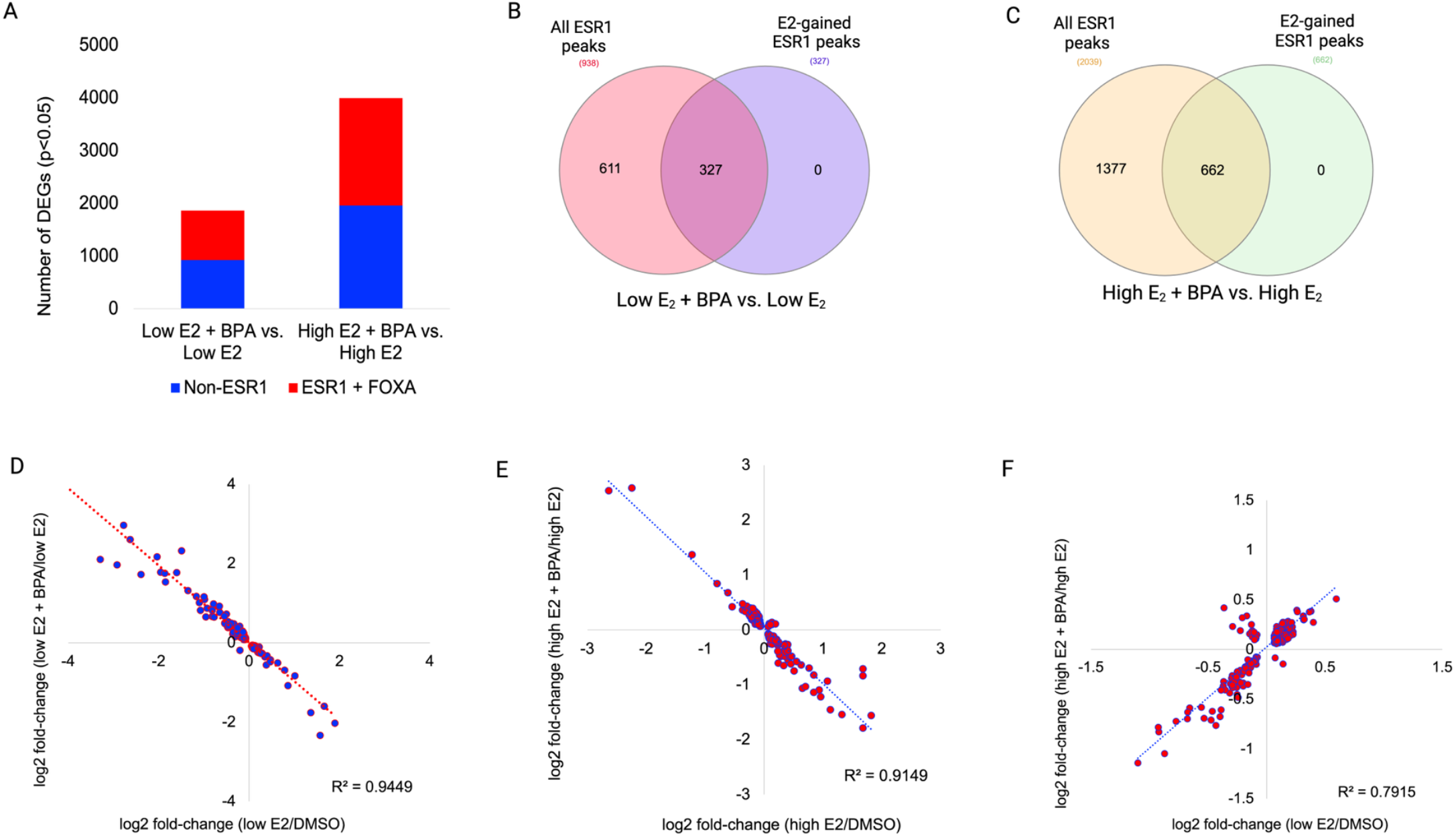
BPA exposure disrupts protective ESR1-mediated gene expression in HepG2 cells. **A**. Barcharts of differentially expressed genes (DEGs) with ESR1 ChIP-seq peak within 50kb and FOXA1/2 motif within 10kb in HepG2 cells exposed to either low E_2_ (5 pM) and BPA (0.4 μM) vs. low E_2_ alone or high E_2_ (10 nM) and BPA vs. high E_2_ alone (*N*=3 per group). **B-C**. Venn diagrams showing overlap of all DEGs near ESR1 ChIP-seq peaks with DEGs near ESR1 ChIP-seq peaks that were newly gained in the presence of E_2_ in healthy human hepatocytes^1^. **D-F**. Scatterplots comparing log_2_ fold-change of DEGs between two comparisons groups (restricted to DEGs p<0.05 present in both groups for each comparison): (D) low E_2_ + BPA vs. low E_2_ compared to low E_2_ vs. DMSO; (E) high E_2_ + BPA vs. high E_2_ compared to high E_2_ vs. DMSO; and (F) high E_2_ + BPA vs. high E_2_ compared to low E_2_ vs. DMSO.

Next, we performed GO enrichment for filtered gene subsets in both comparisons. Both show overrepresentation of terms linked to carcinogenesis (**Supp. Fig. 5**), including “phenylalanine, tyrosine and tryptophan biosynthesis”, “chemical carcinogenesis”, and “non-alcoholic fatty liver disease”, which further validates their functional relevance to our observed cancer phenotype. When we compared expression of the filtered subset of genes in the low E_2_ + BPA vs. low E_2_ and low E_2_ vs. DMSO groups (limited to DEGs present in both comparison groups), we observed a negative slope, indicating that the addition of BPA to the culture reversed the transcriptional effects of low E_2_ on these genes (**Fig. 4D**). This result is consistent with BPA blocking expression changes resulting from low E_2_ alone. We anticipated that we would not observe this result in the high E_2_ + BPA vs. high E_2_ group, as compared to high E_2_ vs. DMSO; surprisingly, we observed a similar pattern in this comparison (**Fig. 4E**). However, when we compared the high E_2_ + BPA group vs. high E_2_ to the low E_2_ vs. DMSO group, we observed a positive slope (**Fig. 4F**), indicating that the addition of BPA to the culture in the presence of high E_2_ yielded similar expression outcomes as did low E_2_ alone. Taken together, these data indicate that BPA is not wholly outcompeted by post-pubertal levels of E_2_ and that co-exposure results in attenuated but not abolished ESR1 target gene expression. Therefore, our data supports the pre-pubertal developmental window as uniquely vulnerable to BPA-mediated disruption of ESR1 target gene expression but suggests that a moderated vulnerability persists into adulthood. Overall, these results are consistent with our model that developmental exposure to BPA promotes liver cancer by via ESR1 binding near FOXA sites and blocking target gene expression in the presence of low (female pre-pubertal) but not high (female post-pubertal) E_2._

## IV. Discussion

Increasing evidence suggests that BPA, a model EDC, is carcinogenic, but mechanistic support is limited^3-7^. We have previously reported that C57BL/6J mice exposed during development to BPA showed linear increases in HCC incidence in adulthood^4^. Here, we provide *in vitro* evidence supporting a mechanism for BPA as a cancer promoting agent a human liver cancer cell line. However, our data do not support BPA as a mutagen at low doses. These data provide the first mechanistic evidence supporting BPA as a non-genotoxic carcinogen in liver. Multiple classes of EDCs, including phenols, phthalates and PFAS, are weak ESR1 agonists^53^ and are associated with adverse liver outcomes in mice and humans^54^, indicating that our proposed mechanism may apply to any EDCs that are weak ESR1 agonists. Therefore, our findings suggest that EDCs should be evaluated more broadly as potential liver carcinogens.

Using a highly sensitive, error-corrected sequencing method, we demonstrate that low doses of BPA, equivalent to human population exposures, are sufficient to cause mutations consistent with oxidative DNA damage but are not sufficient to induce a mutation load greater than background. However, these results do not rule out the possibility that low dose BPA exposure during development increases mutation load *in vivo*. We note that we detected a clear, if muted, mutation signal despite the constraints in our experimental design, including a short exposure duration (24 hours) in a cancer cell line with high mutation background. Notably, mutation spectra previously reported for BPA are highly correlated with the mutation spectra seen in human liver cancer samples from established tissue banks^32^, supporting a possible role for BPA exposure in liver cancers in human populations. Therefore, our results provide sufficient justification for future studies that test the capacity of chronic, low dose BPA exposure during development to initiate liver cancer *in vivo*. These studies will mimic human population exposure scenarios more closely than our *in vitro* experiments. True mutation load in humans exposed during early life is likely to be higher than we observed, because human exposure to BPA is chronic and DNA repair capacity is lower during fetal and infant development, as compared to adulthood.

Our results support early life development as a window of vulnerability to liver cancer promotion by BPA. We show that, in the presence of pre-pubertal, but not post-pubertal, levels of E_2,_ BPA inhibits transcription of genes near ESR1 binding sites that co-localize with FOXA1/2 motifs in healthy human liver. These findings are consistent with our proposed mechanism of HCC promotion by BPA in early life. If this mechanism is confirmed *in vivo*, then BPA is a non-genotoxic carcinogen in liver. This finding is critically important to prevention of HCC, which is prevalent, lethal, and poorly responsive to therapy.

HCC is the fifth most common cancer with the third highest mortality rate worldwide^20,21,55^. This form of cancer has limited treatment options and carries a poor prognosis (17% 5-year survival rate)^20,21,55^. Known HCC risk factors include hepatitis B or C infection, alcohol abuse, or environmental contaminants like aflatoxin B_1_ (AFB_1_)^20^. These risk factors are most common in East Asia, South America, and Africa, which have proportionally higher HCC rates^20^. However, HCC incidence and mortality in Western countries are increasing^20^; liver cancer is poised to become the third leading cause of cancer-related death in the U.S. by 2030^56^. New risk factors, including non-alcoholic steatohepatitis, account for less than a fifth of U.S. cases^20^, suggesting the existence of unidentified environmental risk factors. EDCs, including BPA, have steadily increased chemical number and affected consumer products since the 1970s. Our results indicate that evaluation of EDCs as contributors to HCC risk is strongly warranted.

## V. Methods

### Cell culture and inductions

We grew HepG2 cells (ATCC HB-8065) in T75 flasks in DMEM supplemented with 10% fetal bovine serum (HyClone). Cells were authenticated using short tandem repeat analysis with the GenePrint 10 kit (Promega) and cells tested negative for mycoplasma with the ABM Mycoplasma PCR Detection Kit (catalog no. G238).

For mutation experiments, we used a single T75 flask to seed 12 wells in six-well plates with 0.3 x 10^6^ cells and allowed cells to adhere to wells for 24h. We then added fresh media to three groups of four wells each for 24h: a) 0.4 μM BPA, b) 0.04 μM BPA, or c) DMSO vehicle control (final DMSO concentration 0.5% or less). These doses of BPA are the *in vitro* liver equivalents of human exposure to 5 μg/kg BW/day or 0.5 μg /kg BW/day, respectively, as calculated using the National Toxicology Program’s PBPK tool in their Integrated Chemical Environment. Following exposure, we allowed cells to replicate for 48h. We harvested cells with 0.25% trypsin, pelleted cells by centrifugation for 3 min at 500g, washed the pellets twice with 1X PBS, and extracted DNA using Qiagen DNeasy Blood & Tissue Kit (catalog no. 69504).

For RNA-seq experiments, we switched three T75 flasks of cells to DMEM containing 10% charcoal-stripped/Dextran-treated fetal bovine serum (HyClone) for five days prior to BPA exposure. We grew flasks with continual exposure to E_2_ (10 nM, 5 pM, or DMSO vehicle control) during this period, adding fresh media daily. On Day 6, we seeded 24 wells in six-well plates with 0.3 x 10^6^ cells and allowed cells to adhere to wells for 24h in their respective E_2_ media. On Day 8, we added fresh media to six groups of three wells each for 24h: 1) 0.4 μM BPA; 2) 5 pM E_2_ (equivalent to circulating levels in pre-pubertal females); 3) 10 nM E_2_ (equivalent to circulating levels in post-pubertal females); 4) 0.4 μM BPA + 5 pM E_2_ (modeling an exposed pre-pubertal female); 5) 0.4 μM BPA + 10 nM E_2_ (modeling an exposed post-pubertal female); or 6) DMSO vehicle control. We harvested cells with 0.25% trypsin, pelleted cells by centrifugation for 3 min at 500g, washed pellets with 1X PBS, flash froze in liquid nitrogen and stored immediately at −80°C. We extracted RNA using Qiagen QIAzol Lysis Reagent (catalog number 79306) and Qiagen miRNEasy Mini Kit (catalog number 217004) with additional DNase treatment using Qiagen RNase-free DNase Set (catalog number 79254). Samples were stored immediately at −80°C.

### Duplex sequencing and analysis

We performed error-corrected Duplex sequencing on 12 samples (two exposure groups and one control group, four replicates per group), using the TwinStrand Biosciences Duplex Sequencing mutagenesis kit, as previously described^13,36,57^. This kit uses a 48 Kb panel targeting 20 arbitrary regions throughout the genome. These regions are balanced to provide unbiased sampling of representative sequence contexts, considering GC content, genic/non-genic sequence, coding/non-coding sequence, and other factors; these sites have no known role in cancer and are unlikely to be under selective pressure. Briefly, we sheared double-stranded DNA and ligated fragments with double-stranded unique molecular identifiers (UMIs). Following ligation, the individually labeled strands were PCR amplified, resulting in many copies with shared UMI sequence. After sequencing, we analyzed the resulting dataset using a custom bioinformatic pipeline: https://github.com/Kennedy-Lab-UW/Duplex-Seq-Pipeline. Briefly, we grouped reads sharing the same UMI and computed a consensus sequence for each position in the read for each family to create a “single-strand consensus sequence (SSCS)”. Therefore, each SSCS represented an individual strand of DNA. The SSCS cannot filter sequence errors due to PCR; to filter these, the complementary UMI derived from the same duplex DNA among SSCS reads were compared and the base at each position was retained in the final consensus only if the two strands matched perfectly at that position. We filtered apparent mutations occurring in only one SSCS of each duplex paired consensus and mapped these “duplex consensus sequences (DCS)” back to the reference genome. Any residual deviations from the reference genome were called as true mutations.

### RNA-seq and analysis

We performed stranded RNA-seq on 24 samples (six exposure groups, three replicates per group.) We depleted ribosomal RNA using the Illumina RiboGold kit, generated libraries using Illumina Stranded Total RNA kits, and sequenced barcoded, pooled libraries on an Illumina NovaSeq 6000 S4 flow cell. We evaluated read quality using FastQC and trimmed reads to remove adapters and low-quality reads with Trimmomatic. We aligned trimmed reads to the human reference genome from Ensembl (GRCh38) with STAR and obtained gene counts from the STAR ReadsPerGene files produced according to the Ensembl GRCh38.89 annotation gtf file. For each replicate, we obtained a minimum of 49,689,366 total gene counts. We used edgeR (v3.28.0) to perform differential expression analysis between our groups of interest. We filtered differentially expressed genes based on proximity to an ESR1 site (transcription start site within 50Kb of an ESR1 binding peak from published ChIP data in healthy human liver^1^). Genes passing this filter were further filtered for those with nearby ESR1 peaks with FOXA1 or FOXA2 motif within 10Kb of the ESR1 peak center. We performed GO term enrichment with ShinyGO v0.741 https://bioinformatics.sdstate.edu/go74/

## Funding and Acknowledgements

C.W. acknowledges support from the Oregon Institute of Occupational Health Sciences at Oregon Health & Science University (OHSU) via funds from the Division of Consumer and Business Services of the State of Oregon (ORS 656.630), as well as from an OHSU Center for Developmental Health Seed Grant and an NIH/NIEHS-funded R01 (R01ES034836). Short read sequencing assays were performed by the OHSU Massively Parallel Sequencing Shared Resource. Bioinformatic analysis was performed by the OHSU Epigenetics Consortium. Computational resources and data storage were provided by the OHSU Advanced Computing Center and Exacloud Cluster.

## Author contributions

CW designed and supervised experiments and wrote the manuscript. EW performed cell culture experiments. SC performed DNA and RNA extractions. SK generated Duplex-seq libraries and conducted bioinformatic analysis on Duplex-seq datasets. RS generated RNA-seq libraries, sequenced Duplex-seq and RNA-seq libraries, and aligned all datasets to the human genome. BD and LC performed bioinformatic analyses on RNA-seq datasets. MT advised on cell culture experiments. CA and RSL advised on experimental design. All authors provided critical feedback on the manuscript.

## Competing Interests

The authors declare no competing interests.

## Notes

### Competing Interest Statement

The authors have declared no competing interest.

