## Supplemental Figures for "*In vitro* evidence for bisphenol A as a human liver carcinogen: Environmentally relevant doses inhibit cancer-protective ESR1 signaling in a human liver cell line"

Caren Weinhouse

**This PDF file includes:**

Figures S1 to S5

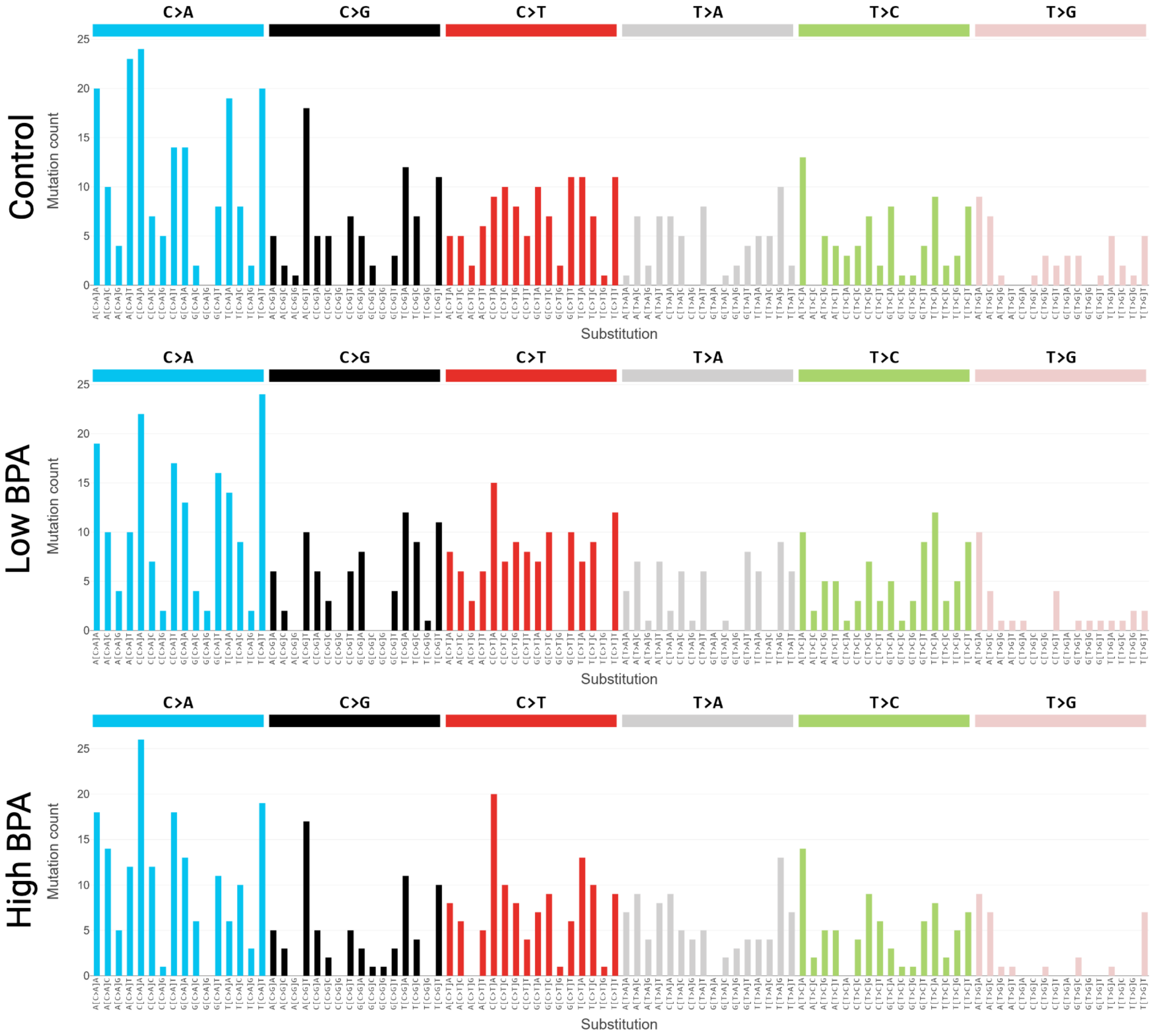


**Supplementary Figure S1. COSMIC Single-Base Substitution (SBS) spectra in BPA-exposed HepG2 cells.** COSMIC SBS spectra in human hepatocellular carcinoma HepG2 cells exposed in triplicate for 24 hours to one of two doses of bisphenol A (BPA) (0.4 µM, or high BPA, or 0.04 µM, or low BPA) or DMSO vehicle control. Cells were cultured for 48 hours post-exposure, to allow for mutation fixation, followed by error-corrected Duplex sequencing of 48 Kb of DNA across 20 genomic regions to ~15,000X and compared mutation frequencies using a custom bioinformatics pipeline (see Methods).

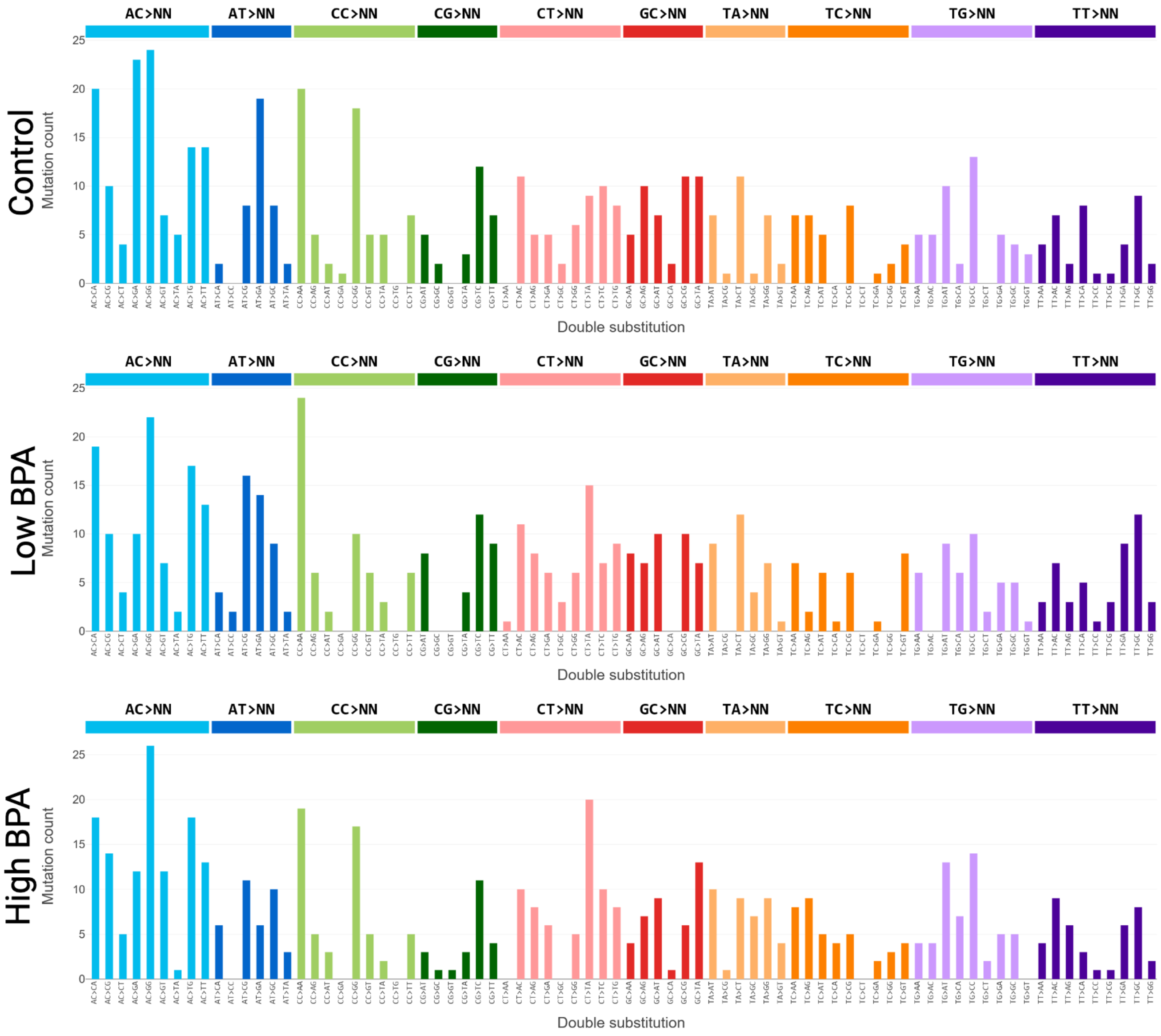


**Supplementary Figure S2. COSMIC Double-Base Substitution (DBS) spectra in BPA-exposed HepG2 cells.** COSMIC DBS spectra in human hepatocellular carcinoma HepG2 cells exposed in triplicate for 24 hours to one of two doses of bisphenol A (BPA) (0.4 $\mu$M, or high BPA, or 0.04 $\mu$M, or low BPA) or DMSO vehicle control. Cells were cultured for 48 hours post-exposure, to allow for mutation fixation, followed by error-corrected Duplex sequencing of 48 Kb of DNA across 20 genomic regions to ~15,000X and compared mutation frequencies using a custom bioinformatics pipeline (see Methods).

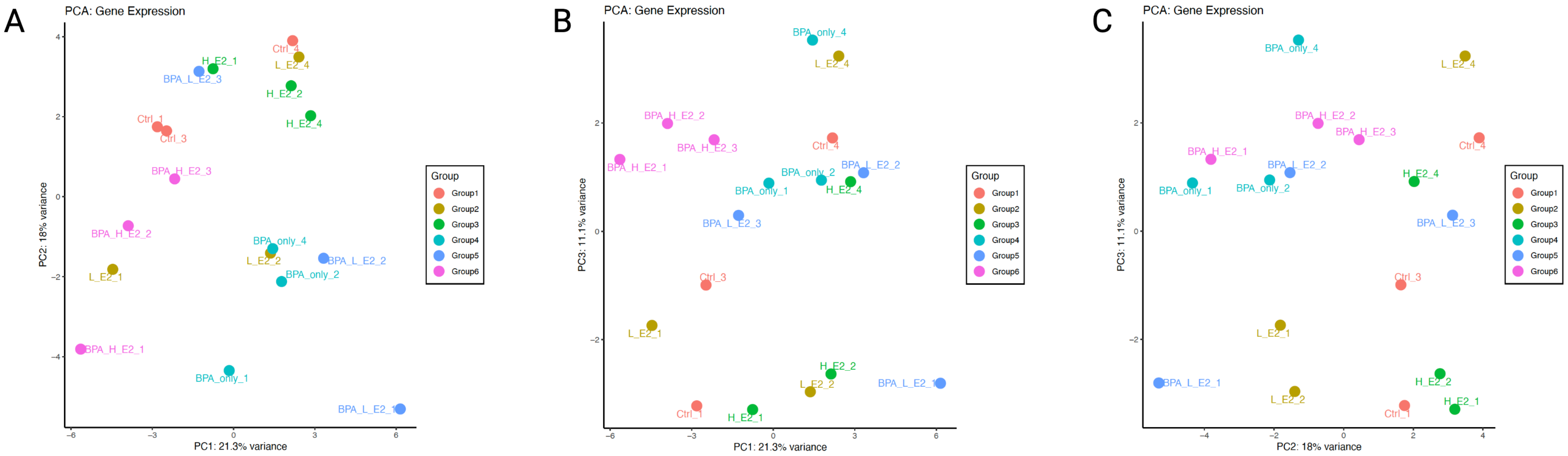


**Supplementary Figure S3. Principal Components Analysis (PCA) plots for RNA-seq datasets in BPA-exposed HepG2 cells.** PCA plots for the first three principal components of RNA-seq datasets generated in HepG2 cells exposed to one of six exposures for 24 hours, following five-day pre-incubation in charcoal-stripped fetal bovine serum and either vehicle (DMSO), low $E_2$, or high $E_2$, to mimic the group-relevant background hormonal environment: 1) 0.4 $\mu$M BPA; 2) 5 pM $E_2$ (equivalent to circulating levels in pre-pubertal females); 3) 10 nM $E_2$ (equivalent to circulating levels in post-pubertal females); 4) 0.4 $\mu$M BPA + 5 pM $E_2$ (modeling an exposed pre-pubertal female); 5) 0.4 $\mu$M BPA + 10 nM $E_2$ (modeling an exposed post-pubertal female); or 6) DMSO vehicle control. (A) Plot of PCA components 1 and 2. (B) Plot of PCA components 1 and 3. (C) Plot of PCA components 2 and 3.

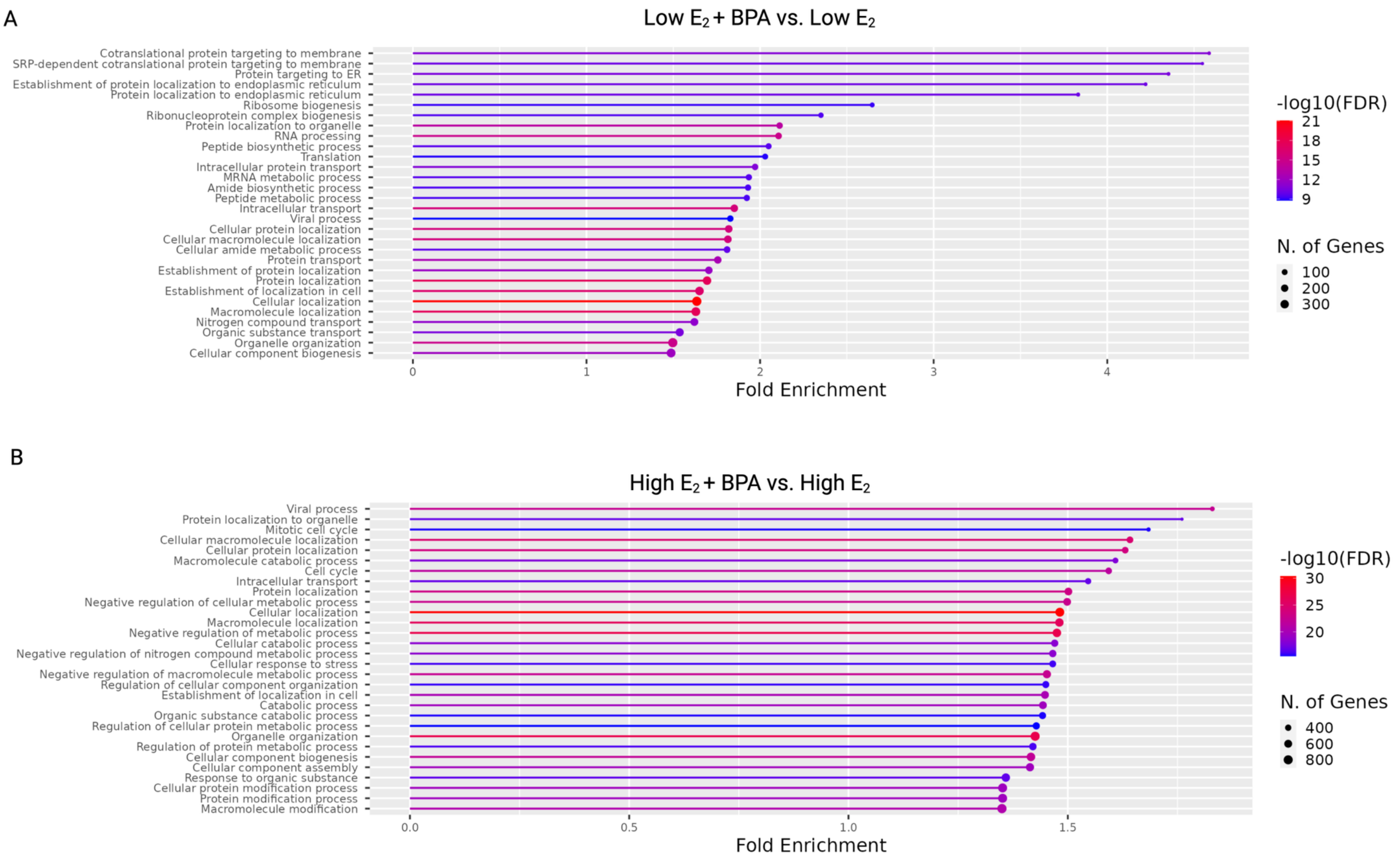


**Supplementary Figure S4. Gene Ontology (GO) enrichment in differentially expressed genes in BPA-exposed HepG2 cells.** GO enrichments for differentially expressed genes in HepG2 cells in one of two comparison groups: (A) Low $E_2$ + BPA vs. Low $E_2$ or (B) High $E_2$ + BPA vs High $E_2$. All exposures were for 24 hours, following five-day pre-incubation in charcoal-stripped fetal bovine serum and either vehicle (DMSO), low $E_2$, or high $E_2$, to mimic the group-relevant background hormonal environment. Cells were exposed to one of six exposures: 1) 0.4 $\mu$M BPA; 2) 5 pM $E_2$ (equivalent to circulating levels in pre-pubertal females); 3) 10 nM $E_2$ (equivalent to circulating levels in post-pubertal females); 4) 0.4 $\mu$M BPA + 5 pM $E_2$ (modeling an exposed pre-pubertal female); 5) 0.4 $\mu$M BPA + 10 nM $E_2$ (modeling an exposed post-pubertal female); or 6) DMSO vehicle control. (A) shows data comparing groups (4) and (2). (B) shows data comparing groups (5) and (3).

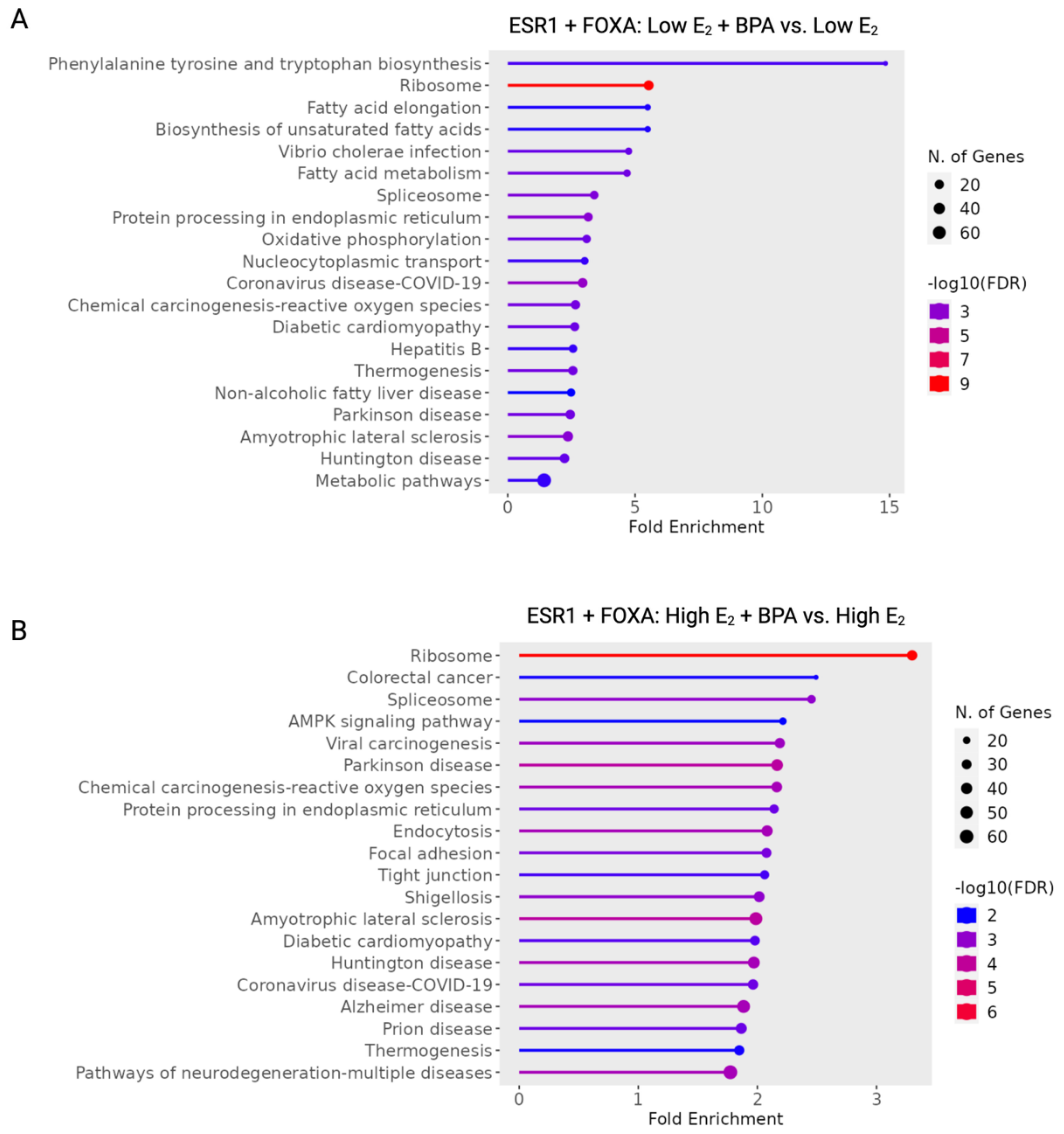


**Supplementary Figure S5. Gene Ontology (GO) enrichment in differentially expressed ESR1 target genes in BPA-exposed HepG2 cells.** GO enrichments for differentially expressed genes in HepG2 cells, filtered for genes within 50 Kb of a published ESR1 ChIP-seq peak and within 10 Kb of a FOXA1/2 motif, in one of two comparison groups: (A) Low $E_2$ + BPA vs. Low $E_2$ or (B) High $E_2$ + BPA vs High $E_2$. All exposures were for 24 hours, following five-day pre-incubation in charcoal-stripped fetal bovine serum and either vehicle (DMSO), low $E_2$, or high $E_2$, to mimic the group-relevant background hormonal environment. Cells were exposed to one of six exposures: 1) 0.4 μM BPA; 2) 5 pM $E_2$ (equivalent to circulating levels in pre-pubertal females); 3) 10 nM $E_2$ (equivalent to circulating levels in post-pubertal females); 4) 0.4 μM BPA + 5 pM $E_2$ (modeling an exposed pre-pubertal female); 5) 0.4 μM BPA + 10 nM $E_2$ (modeling an exposed post-pubertal female); or 6) DMSO vehicle control. (A) shows data comparing groups (4) and (2). (B) shows data comparing groups (5) and (3).
